# The fungal sesquiterpenoid pyrenophoric acid B uses the plant ABA biosynthetic pathway to inhibit seed germination

**DOI:** 10.1101/622530

**Authors:** Jorge Lozano-Juste, Marco Masi, Alessio Cimmino, Suzette Clement, Maria A. Fernández, Regina Antoni, Susan Meyer, Pedro L. Rodriguez, Antonio Evidente

## Abstract

The fungus *Pyrenophora semeniperda* produces pyrenophoric acid B, a small molecule that exploits the plant ABA biosynthetic pathway to reduce seed germination, increasing its reproductive success.

**Abstract:** Pyrenophoric acid (P-Acid), P-Acid B and P-Acid C are three phytotoxic sesquiterpenoids produced by the ascomycete seed pathogen *Pyrenophora semeniperda,* a fungus proposed as a mycoherbicide for biocontrol of cheatgrass, an extremely invasive weed. When tested in cheatgrass bioassays these metabolites were able to delay seed germination, with P-Acid B being the most active compound. Here, we have investigated the cross-kingdom activity of P-Acid B and its mode of action and found that it activates the ABA signaling pathway in order to inhibit seedling establishment. P-Acid B inhibits seedling establishment in wild-type *Arabidopsis thaliana* while several mutants affected in the early perception as well as in downstream ABA signaling components were insensitive to the fungal compound. However, in spite of structural similarities between ABA and P-Acid B, the latter is not able to activate the PYR/PYL family of ABA receptors. Instead, we have found that P-Acid B uses the ABA biosynthesis pathway at the level of alcohol dehydrogenase ABA2 to reduce seedling establishment. We propose that the fungus *Pyrenophora semeniperda* manipulates plant ABA biosynthesis as a strategy to reduce seed germination, increasing its ability to cause seed mortality and thereby increase its fitness through higher reproductive success.

## Introduction

The fungus *Pyrenophora semeniperda* (Brittlebank and Adams) Shoemaker, a naturally occurring necrotrophic seed pathogen, has been proposed as a potential biocontrol agent against cheatgrass *(Bromus tectorum)* (Meyer *et al.*, 2013). This invasive weed is dramatically altering the semi-arid shrubland ecosystems in the western U.S.A. increasing wildfire frequency and intensity (Germino *et al.*, 2016). Thus, the ability of *P. semeniperda* to produce toxins *in vitro*, which could potentially be useful in biological control, was investigated (Meyer *et al.*, 2015; Cimmino *et al.*, 2015). *P. semeniperda* produces large amounts of cytochalasin B in solid cultures (Masi *et al.*, 2014a) while spirocyclic γ-lactams, including spirostaphylotrichins A, C, D, R, V, W and triticone E, are produced when the fungus is grown in potato dextrose broth (Masi *et al.*, 2014b). Furthermore, from the same fungal organic extract smaller quantities of cytochalasins A, F and deoxaphomin were isolated together with a new phytotoxic sesquiterpenoid, named pyrenophoric acid (P-Acid) (Fig. 1A) (Masi *et al.*, 2014a,c). Finally, these compounds were produced in *B. tectorum* seed culture together with other two phytotoxic sesquiterpenoids named pyrenophoric acids B and C (P-Acid B and P-Acid C, respectively) (Fig. 1A) (Masi *et al.*, 2014d). P-Acid, P-Acid B and P-Acid C were able to inhibit both root and coleoptile growth of *B. tectorum* (Masi *et al.*, 2014c,d). In particular, P-Acid B was the most active sesquiterpenoid among the P-Acids studied (Masi *et al.*, 2014d). It was able to reduce germination at 7 days after sowing and produced germination delay relative to the control treatment. P-Acid B was also more bioactive in reducing coleoptile and root growth than the other sesquiterpenoids, P-Acid and P-Acid C. Interestingly, the production of small molecules able to delay germination could benefit *P. semeniperda*, as its fitness is increased on slow-germinating or dormant seeds. This is because slow-germinating seeds cannot escape pathogen-caused mortality, thus increasing resources available for pathogen growth and reproduction (Meyer *et al.*, 2015; Cimmino *et al.*, 2015).

**Figure 1.**
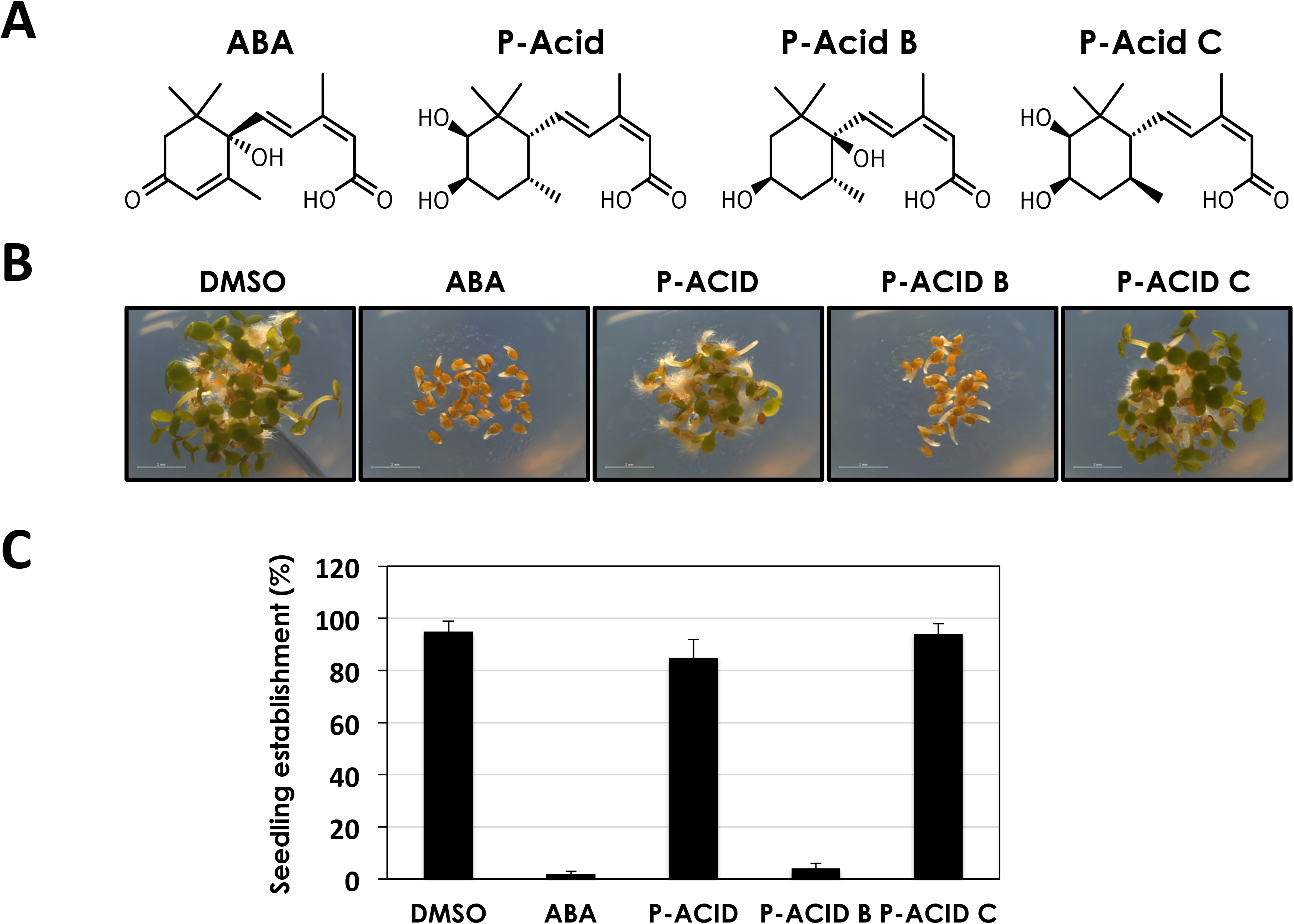
Pyrenophoric acid B inhibits seed germination. (A) Structures of abscisic acid, ABA, pyrenophoric acids, P-Acid, P-Acid B and P-Acid C used in this work. (B) Pictures of Arabidopsis Col-0 seeds treated with DMSO, 1 μM ABA, and 100 μM of the different pyrenophoric acids, 3 days after shown. (C) Quantification of seedling establishment 3 days after shown Col-0 seeds on DMSO, 1 μM ABA, and 100 μM of the different pyrenophoric acids. Values represent mean ± SD of 90 seed used in three different experiments.

P-Acid, P-Acid B and P-Acid C (Fig. 1A) are sesquiterpenoids closely related to abscisic acid (ABA) (Fig. 1A). ABA is a plant hormone involved in many physiological processes in plants including seed development and germination, root growth and abscission among others (Leung and Giraudat, 1998). It has also a central role in the adaptation to environmental stresses like drought, high salinity and low temperatures (Yamaguchi-Shinozaki and Shinozaki, 2006). However, ABA is not a plant-specific metabolite. It is produced by different organisms including fungi, suggesting that it is a ubiquitous and versatile small compound that can modulate a myriad of physiological functions (Hartung, 2010; Takezawa *et al.*, 2011). In *Arabidopsis thaliana* (Arabidopsis), ABA biosynthesis begins with the conversion of zeaxanthin to all-trans-violaxanthin by a zeaxanthin epoxidase known as ABA1 (Marin et al., 1996, Audran et al., 2001). Next, in the rate-limiting step of ABA biosynthesis, 9-cis-epoxycarotenoid dioxygenase (NCED) enzymes break down C40 carotenoids to produce xanthoxin (Schwartz *et al.*, 1997; Iuchi *et al.*, 2000). Xanthoxin is further transformed into abscisic aldehyde by the Arabidopsis short-chain alcohol dehydrogenase ABA2 (González-Guzmán *et al.*, 2002). Finally, abscisic aldehyde is used as substrate by the aldehyde oxidase AAO3, in a reaction that depends on the activity of the molybdenum cofactor sulfurase ABA3, to produce ABA (Schwartz *et al.*, 1997; Seo *et al.*, 2000; González-Guzmán *et al.*, 2004). ABA is perceived by the PYR/PYL/RCAR family of ABA receptors, comprised of 14 members in the eudicot plants Arabidopsis and tomato and 12 and 9 members in the monocot species rice and *Brachypodium distachyon,* respectively (Ma *et al.*, 2009; Park *et al.*, 2009; He *et al.*, 2014; Pri-Tal *et al.*, 2017). ABA-bound PYR/PYL/RCAR proteins interact with Protein Phosphatases type 2C (PP2C) and inhibit their activity allowing the downstream phosphorylation cascades to activate the ABA response, including inhibition of seed germination and stomatal closure, among others.

Here we have studied the effect and the mode of action of the fungus-produced pyrenophoric acids on plant seed germination. Among P-Acids, P-Acid B was able to inhibit germination of Arabidopsis seeds. We have found that P-Acid B makes use of the ABA biosynthesis pathway at the level of ABA2 to activate the ABA signaling and inhibit germination, increasing the probability of host seed mortality and thereby increasing pathogen fitness.

## Materials and Methods

### Chemicals

Pyrenophoric acid and pyrenophoric acids B and C (Fig. 1A) were isolated from the solid *B. tectorum* (cheatgrass) seed culture of *P. semeniperda* (strain WRK10-22), as previously reported (Masi *et al.*, 2014c,d)

### Plant material and growth conditions

*Arabidopsis thaliana* seeds were surface sterilized with 30% bleach containing 0.02% Tween-20 for 10 minutes followed by rinsing with sterile water 5 times. Seeds were sown on solid or liquid Murashige and Skoog (MS) media and stratified 3 days at 4°C. For seedling establishment experiments, seeds were sown on 24-well plates filled with 1 mL of MS media + 1% agar supplemented with the different chemicals dissolved in DMSO. Seeds were kept in a Sanyo incubator under long day conditions at temperatures 23 °C /21°C (day/night) for 3 days. Pictures were taken with a Leica DMS1000 macroscope and seedling establishment (seedlings with open and green cotyledons) scored. For luciferase assays, seeds were incubated for 7 days under long day conditions in liquid MS media. The different mutants used in this work, *112458* (Gonzalez-Guzman *et al.*, 2012), *abi3-9* (Nambara *et al.*, 2002), *abi4-11* (Nambara *et al.*, 2002), *aba1-101* (Barrero *et al.*, 2005), *aba2-1* (Léon-Kloosterziel *et al.*, 1996), *nced3,5* (Frey *et al.*, 2012) *aba2-11* (González-Guzmán *et al.*, 2002), *aao3-2* (González-Guzmán *et al.*, 2004), *aba3-1* (Léon-Kloosterziel *et al.*, 1996), and the transgenic lines *pMAPKKK18-LUC+* (Vaidya *et al.*, 2017) and *35S-HAB1^W385A^* (Dupeux *et al.*, 2011) have been described previously. To generate the double *nced2nced3* mutant, homozygous plants from the T-DNA lines SALK_090937C *(nced2)* and GK-129B08.01 *(nced3)* were crossed and the F2 selected on 100 mM NaCl plates. PCR-verified plants using the primers FORNced2: ATGGTTTCTCTTCTTACAATGCCG, RVNced2: TTCCGGTTAACCATACCAATCTC, NewpROK2: GCCGATTTCGGAACCACCATC, FWNced3GABI: CCTAGTGTTCAGATCGCCGGA, RVNced3GABI: GAGATTCTCGTCAGACTCGTT, GK-o8474: ATAATAACGCTGCGGACATCTACATTT were selfed and the F3 seed was used for the different experiments.

### Luciferase imaging

*pMAPKKK18-LUC*^+^ seedlings were grown in 24-well plates filled with 1 mL liquid MS media for 7 days. MS media was changed with MS media supplemented with 100 μM D-luciferin, potassium salt (GoldBio) and the different treatments; 25 μM ABA, 200 μM P-Acid B or DMSO as control, and incubated for 6 hours. Luminescence was recorded with a LAS-3000 imager (Fujifilm) equipped with a CCD camera using 2 minutes exposures. 8-bit images were transformed in rainbow false color and quantified using Fiji (Schindelin *et al.*, 2012).

### Protein purification

The different expression clones were described previously (Antoni *et al.*, 2012; Castillo *et al.*, 2015). Protein induction was performed by adding 1mM IPTG to 250mL cultures of BL21(DE3) cells at OD=0.6-0.8. Cells were grown overnight at 16 ^o^C and collected by centrifugation. Cells were lysed in 50 mM tris-HCl (pH 7.6), 250 mM KCl, 10mM 2-ME, 10mM Imidazole, 0.1% Tween 20, 10% glycerol and freeze/thawed at −80°C twice. After sonication, the protein extract was cleared by centrifugation and applied into 1mL Ni-NTA agarose. Ni-NTA beads were washed 3 times with 10 column volumes of 50 mM tris-HCl (pH 7.6), 250 mM KCl, 10mM 2-ME, 20% glycerol, 0.1% Tween 20 buffer supplemented with 30mM imidazole. Proteins were eluted from the Ni-NTA beads with the same buffer containing 250mM imidazole.

### PP2C enzymatic assays

For PP2C assays 1μM dNHAB1 and 2 μM PYR/PYL were used (Santiago *et al.*, 2009). Proteins were incubated with the different chemicals (DMSO, ABA or P-Acid B) for 10 minutes in a volume of 50 μL containing 10mM MnCl_2_. Reactions were initiated by the addition of 50 μL 50mM Tris pH=8.0, 50mM p-nitrophenyl phosphate (pNPP). Dephosphorylation of pNPP was followed by monitoring the absorbance at 405 nm during 20 minutes. The activity of dNHAB1 in the absence of any receptor was set to 100%.

## Results

### Effect of pyrenophoric acids on seed germination

P-Acid, P-Acid B and C (Fig. 1A) were purified from an organic extract of *P. semeniperda* (strain WRK10-22) grown in solid *B. tectorum* (cheatgrass) seed culture following the procedure described previously (Masi *et al.*, 2014c,d) and identified by comparing the physical and spectroscopic properties with those previously reported (Masi *et al.*, 2014c,d). To assess the effects of P-Acids *in planta,* Arabidopsis WT Col-0 seeds were germinated in the presence of 100 μM of each P-Acid, 1 μM ABA or DMSO as a control. P-Acid and P-Acid C did not have any effect on seed germination while P-Acid B severely delayed germination and markedly inhibited seedling establishment (Fig. 1B). At day 3 after sowing 100% of Col-0 seedlings established under control (DMSO), P-Acid or P-Acid C conditions. However, the P-Acid B treatment reduced seedling establishment to 10%, similar to a treatment with 1 μM ABA (Fig. 1C). Therefore, it seems that among P-Acids, pyrenophoric acid B, produced by the fungus *P. semeniperda,* has cross-kingdom activity, as it is able to inhibit plant seed germination.

### Pyrenophoric acid B activates the ABA signalling pathway

In order to gain further insight into the mode of action of P-Acid B we used several mutants affected in the early and late steps of ABA perception and signalling. We used the *112458* sextuple mutant impaired in PYR1, PYL1, PYL2, PYL4, PYL5 and PYL8 ABA receptors (Gonzalez-Guzman *et al.*, 2012); 35S:HAB1^W385A^, a transgenic line expressing an engineered mutant form of the PP2C HAB1 unable to interact with ABA receptors (Dupeux *et al.*, 2011); and *abi3* and *abi4* that carry mutations in 2 transcription factors central to the activation of ABA signaling (Nambara *et al.*, 2002). All mutants tested were insensitive to P-Acid B application (Fig. 2A). While only 10% of Col-0 seedlings established in the presence of 100 μM P-Acid B, all the mutants reached nearly 100% seedling establishment 3 days after the treatment (Fig. 2B). These results indicate that the pathogen-produced P-Acid B uses the plant ABA signaling pathway to inhibit seed germination. To elucidate if P-Acid B could activate ABA-dependent gene expression we used an ABA reporter line that carries the ABA inducible promoter *pMAPKKK18* (Okamoto *et al.*, 2013) fused to the firefly luciferase enzyme. This reporter line is highly activated by ABA (Fig. 2C,D) (Vaidya *et al.*, 2017). P-Acid B was able to activate the reporter line since *pMAPKKK18-LUC^+^* seedlings treated with 200 μM P-Acid B during 6 hours showed around 4.5 fold increase on LUC activity due to the activation of the ABA responsive promoter *pMAPKKK18* (Fig. 2D). In conclusion, P-Acid B is able to activate ABA signalling, including ABA-dependent gene expression, to inhibit seed germination.

**Figure 2.**
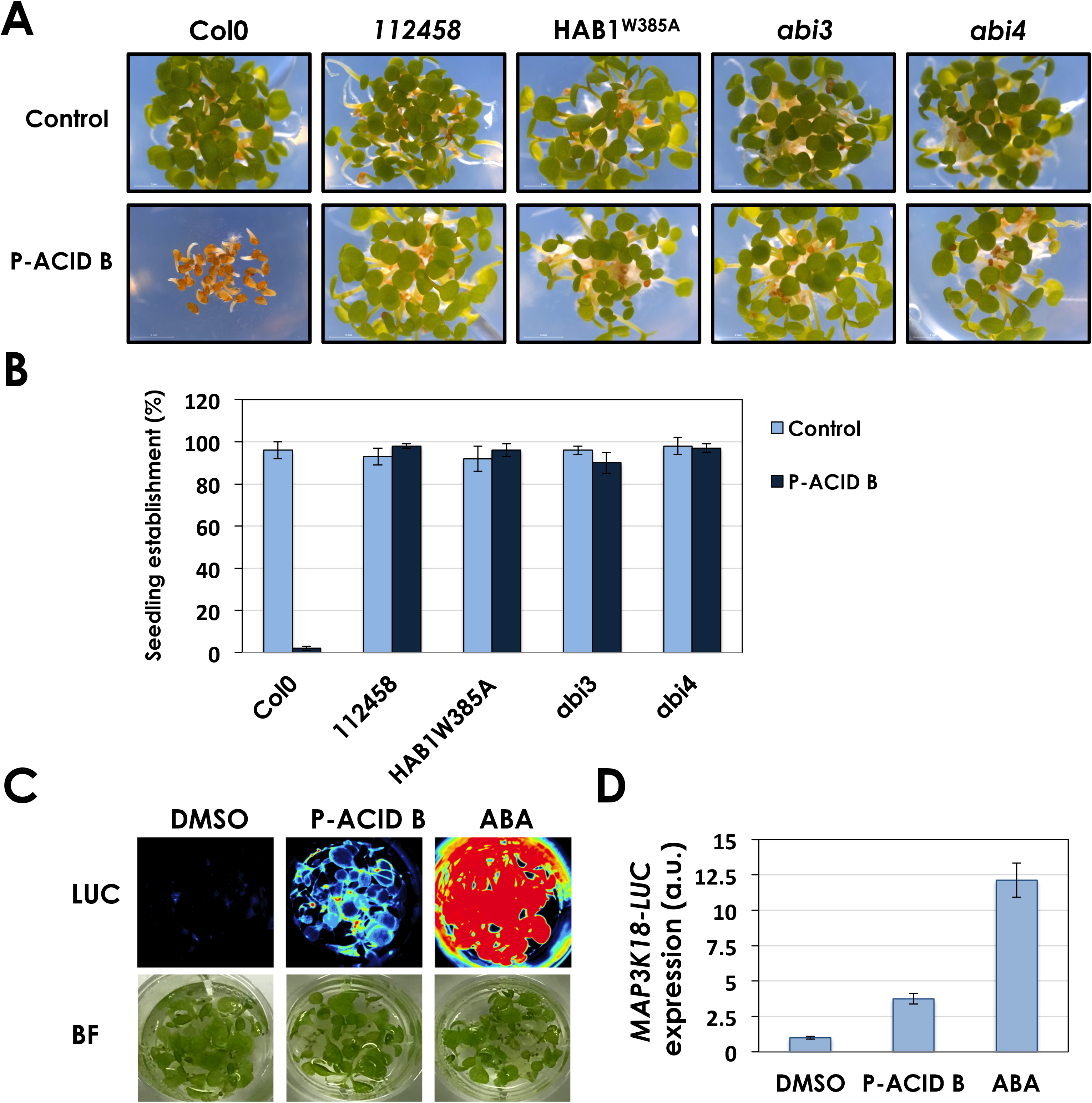
Pyrenophoric acid B (P-Acid B) activates the ABA signaling pathway to inhibit seed germination. (A) Pictures of Arabidopsis seeds treated with 100 μM Pyrenophoric acid B or DMSO as control 3 days after sown. Col-0 seed was used along with mutants in the ABA signaling pathway. (B) Quantification of seedling establishment 3 days after sown seeds on DMSO or 100 μM of P-Acid B. Values represent mean ± SD of 90 seed used in three different experiments. (C) Luminescence detection (LUC) of seedlings of the ABA reporter line pMAPKKK18-LUC+ treated with DMSO as control, 200 μM P-Acid B or 25 μM ABA for 6 hours. The image of the seedlings under bright light (BF) is also included. (D) Quantification of the pMAPKKK18-LUC+ expression shown in C.

### Pyrenophoric acid B does not activate ABA receptors

P-Acid B inhibits seed germination and the sextuple mutant *112458* is insensitive to the application of P-Acid B (Fig. 2A,B). In addition, pyrenophoric acids are sesquiterpenoids with structural similarities to ABA (Fig. 1A). We reasoned that P-Acid B could activate the ABA signaling pathway by directly binding to ABA receptors. To investigate this possibility we used PP2C phosphatase assays with different ABA receptors and the PP2C phosphatase HAB1. The PYR/PYL/RCAR family of ABA receptors binds ABA within a hydrophobic cavity that defines the ABA binding pocket (Santiago *et al.*, 2009; Melcher *et al.*, 2009). After ABA binding, allosteric changes induced in ABA receptors facilitates their interaction with PP2Cs inhibiting phosphatase activity. Therefore, the binding of small molecules to ABA receptors can be followed by *in vitro* PP2C phosphatase assays as reported elsewhere (Park *et al.*, 2009; Okamoto *et al.*, 2013; Vaidya *et al.*, 2017). We performed *in vitro* PP2C phosphatase assays using recombinant PYR1, PYL1, PYL2, PYL4, PYL5, PYL6, PYL8 and PYL10. As expected, ABA bound to all receptors tested producing the concomitant inhibition of HAB1 phosphatase activity by 80-90%. However, P-Acid B was not able to inhibit HAB1 in the presence of any of the receptors tested (Fig. 3), strongly suggesting that despite structural and functional similarities with ABA, P-Acid B does not activate ABA receptors.

**Figure 3.**
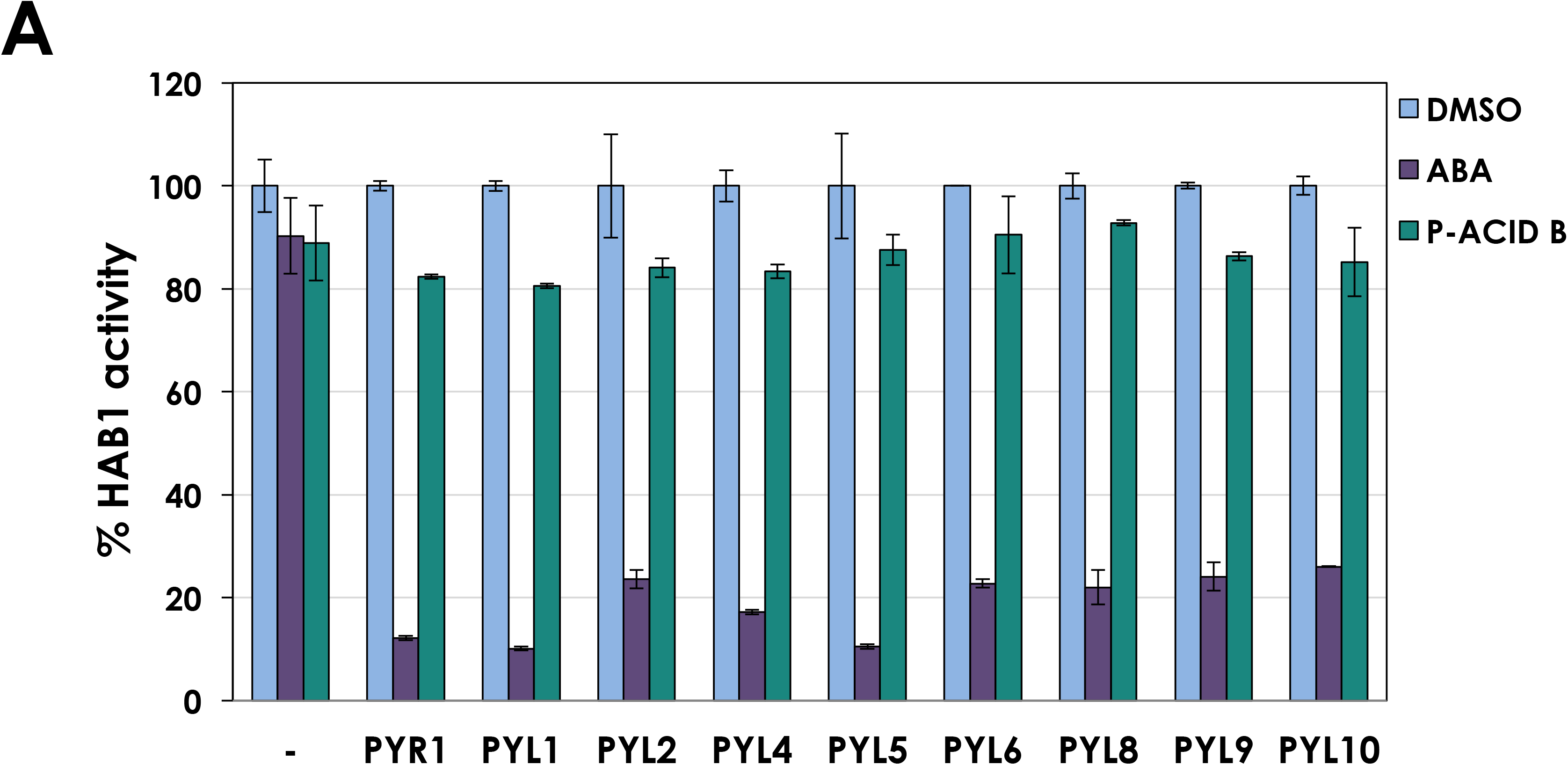
Pyrenophoric acid B (P-Acid B) does not activate ABA receptors. PP2C phosphatase assay using HAB1 (1 μM) and different ABA receptors (2 μM) in the presence of DMSO as control, 10 μM ABA or 100 μM P-Acid B. HAB1 phosphatase activity was set to 100% in the absence of receptor and pNPP was used as a chromogenic substrate. Values represent mean ± SD (n=3).

### P-Acid B exploits the plant ABA biosynthetic pathway to inhibit seed germination

P-Acid B inhibits seed germination and ABA-insensitive mutants are also insensitive to P-Acid B, suggesting that P-Acid B needs an intact ABA signaling pathway to inhibit seed germination. To better understand P-Acid B activity we analyzed the effect of P-Acid B treatment on the germination of mutants affected in different steps of the ABA biosynthetic pathway. We used *aba1-101,* compromised in the early steps of ABA biosynthesis (Barrero *et al.*, 2005), the double mutants *nced2nced3 (nced2,3)* and *nced3,5,* impaired in the production of xanthoxin; *aba2-1* and *aba2-11,* loss of function alleles of *ABA2* that cannot convert xanthoxin into ABA-aldehyde (González-Guzmán *et al.*, 2002; Cheng *et al.*, 2002) and lastly, *aao3-2* and *aba3-1,* blocked in the last step of ABA biosynthesis where ABA-aldehyde is oxidized to ABA (González-Guzmán *et al.*, 2004; Schwartz *et al.*, 1997b). As expected, while all the genotypes germinated and established 3 days after sowing in control (DMSO) conditions, 1 μM ABA treatment inhibited germination and seedling establishment in the wild-type Col-0 and also in all the ABA deficient mutants tested (Fig. 4A,B; Fig. S1). As in previous experiments, P-Acid B delayed germination and severely affected seedling establishment of Col-0 seeds. Interestingly, *aba2-1, aba2-11, aao3-2 and aba3-1* mutants were resistant to P-Acid B treatment indicating that ABA biosynthesis from ABA2 onwards is required for P-Acid B bioactivity on seed germination. However, *nced2,3* which is impaired in the upstream step of ABA biosynthesis that precedes ABA2 activity, was sensitive to P-Acid B, with a clear reduction in seedling establishment in response to this treatment (Fig. 4A,B). This result was confirmed using a different *NCED* double mutant; *nced3,5,* and also the mutant *aba1-101,* impaired in the first step of ABA biosynthesis (Fig. S1). Thus, while P-Acid B reduced seedling establishment of Col-0, *aba1-101, nced2,3* and *nced3,5* seeds to below 10%, the compound was unable to produce any effect on the other ABA biosynthetic mutants *aba2-1, aba2-11, aao3-2* and *aba3-1* (Fig. 4A,B; Fig. S1). Therefore, P-Acid B can complement the ABA biosynthetic defect of *aba1-101, nced2,3* and *nced3,5* but requires downstream enzymatic steps. These results suggest that P-Acid B requires ABA2 activity to inhibit seed germination and establishment. Actually, since P-Acid B is structurally related to xanthoxin and ABA2 is able to convert xanthoxin-related compounds into abscisic aldehyde (González-Guzmán *et al.*, 2002; Nambara and Marion-Poll, 2005), we hypothesized that P-Acid B could be used by ABA2 as a substrate to produce ABA or an ABA-mimic, thus explaining the activation of the ABA signaling pathway by P-Acid B and its negative effect on seed germination.

**Figure 4.**
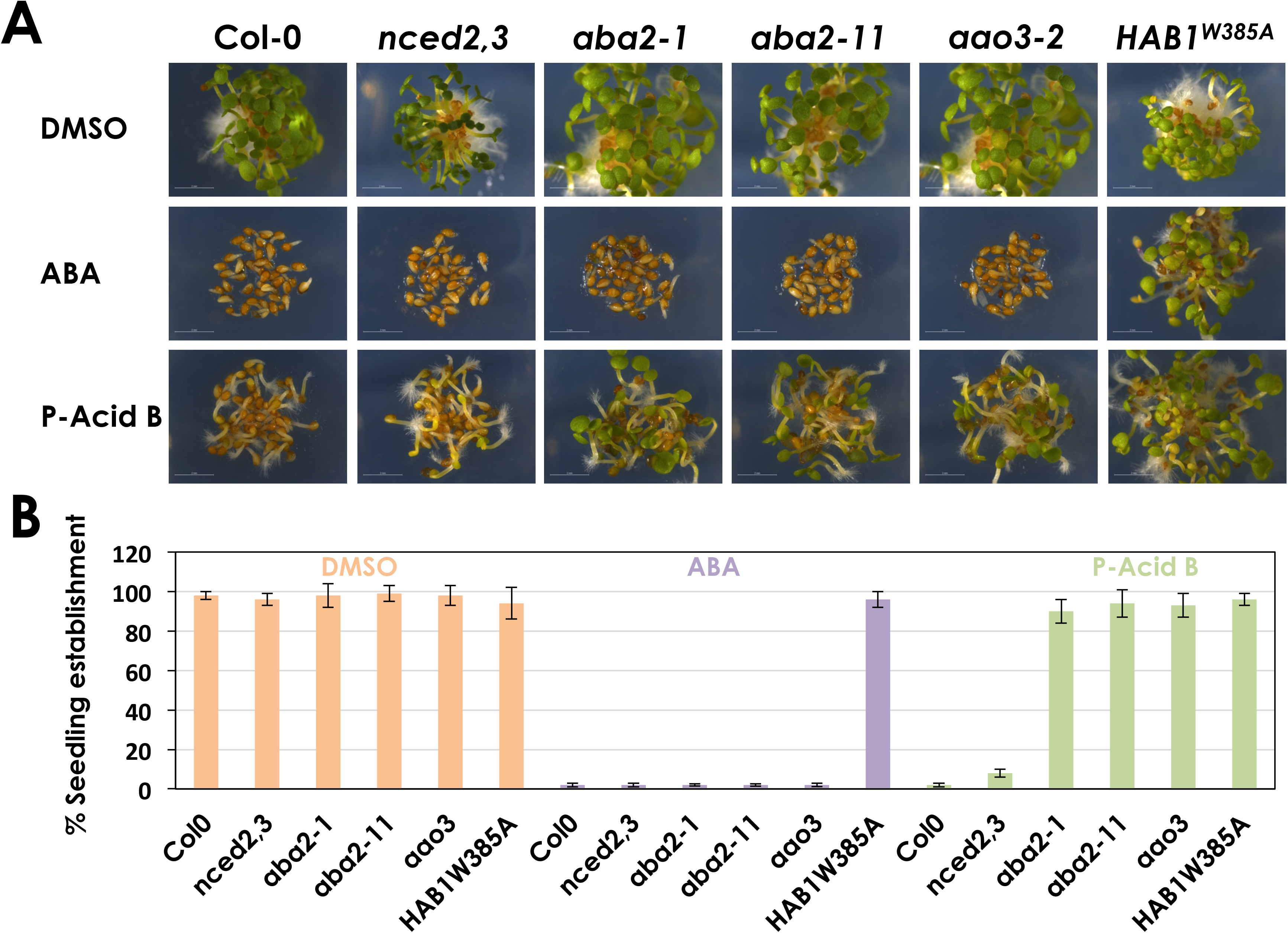
Pyrenophoric acid B (P-Acid B) uses the ABA biosynthesis pathway to inhibit seed germination. (A) Pictures of Arabidopsis seeds treated with 1 μM ABA, 100 μM P-Acid B or DMSO as control, 3 days after sown. Col-0 was used along with mutants in the ABA biosynthesis pathway. (B) Quantification of seedling establishment 3 days after sown seeds on DMSO, 1 μM ABA or 100 μM P-Acid B. Values represent mean ± SD of 90 seed used in three different experiments.

## Discussion

The fungus *P. semeniperda* is a plant pathogen specialized in infecting seed tissues. Due to its ability to infect cheatgrass, one of the most invasive plant species in the USA, it has been lately considered as a biocontrol agent for invasive grass species. For this purpose, *P. semeniperda* produces a series of toxins that kill the host cells, including the mycotoxin cytochalasin B, which impedes the correct polymerization of actin filaments leading to cell death (Scherlach *et al.*, 2010). However, cytochalasin B is not the only low molecular weight compound produced by *P. semeniperda* that has a role in infection. Recently, a series of metabolites was isolated and identified from *P. semeniperda* cultures and among them, P-Acid, P-Acid B and P-Acid C were found (Masi *et al.*, 2014c,d). P-Acids are bioactive compounds able to inhibit seed germination and coleoptile growth, with P-Acid B the most active of these compounds (Masi *et al.*, 2014d). In this work we have investigated the molecular mechanism underlying P-Acids mode of action. We have found that P-Acid B can inhibit *Arabidopsis thaliana* seed germination through the ABA signaling pathway. P-Acid B needs an intact ABA signaling pathway to inhibit seed germination since mutants affected in the first steps of ABA perception as well as mutants impaired in central transcription factors essential for the activation of ABA responses were insensitive to P-Acid B application.

P-Acids are ABA-related compounds with structural similarities to the plant hormone. It has been previously reported that ABA receptors can accommodate ABA-like molecules within its hydrophobic pocket (Kepka *et al.*, 2011; Benson *et al.*, 2015). For example, the ABA degradation by-product, phaseic acid (PA), is able to bind to certain ABA receptors (Weng *et al.*, 2016). PA has submicromolar activity towards the ABA receptors PYL3, PYL5 and PYL6 and a role for PA on water stress tolerance has been proposed (Weng *et al.*, 2016; Lozano-Juste and Cutler, 2016; Rodriguez, 2016). Other hormone receptors can also accommodate hormone analogs or different forms of the hormone. GID1, the gibberellic-acid receptor binds different gibberellins (Murase *et al.*, 2008; Shimada *et al.*, 2008) and the jasmonic acid (JA) receptor COI1 is activated not only by its plant derived ligand, JA-Ile, but also by the JA analog coronatin, produced by biotrophic bacteria (Sheard *et al.*, 2010). Therefore we reasoned that the ABA-related sequiterpenoid P-Acid B could directly bind to ABA receptors. However, we could not find activation to any of the ABA receptors studied indicating that P-Acid B, despite its structural similarities to ABA, does not activate ABA receptors directly, suggesting a different mechanism of action. An alternative hypothesis for P-Acid B bioactivity is acting as precursor of ABA, feeding the biosynthetic pathway and promoting ABA accumulation. This hypothesis proved to be correct since *aba3-1, aao3-2, aba2-1* and *aba2-11* mutants affected in ABA biosynthesis could establish in the presence of P-Acid B while Col-0 plants could not. Therefore, P-Acid B may require *de novo* synthesis of ABA to inhibit seedling establishment. In order to pinpoint the entrance of P-Acid B into the ABA biosynthetic pathway, we have compared sensitivity to P-Acid B of mutants impaired in different steps of ABA biosynthesis. Thus, when we analyzed the effect of P-Acid B on the *aba1-101, nced2,3* and *nced3,5* mutants, we found that these strains are not resistant to P-Acid B, while *aba2*, *aao3* and *aba3* single mutants could germinate and establish in presence of P-Acid B. These data suggest that P-Acid B enters the ABA biosynthetic pathway downstream of NCED and requires ABA2, AAO3 and ABA3 for bioactivity. AAO3 encodes an aldehyde oxidase and might restore the carboxylic group of P-Acid B in case it was reduced to aldehyde by the reductive ability of plant cells for exogenous molecules. Indeed, the reduction of non-activated carboxylic groups in different substrates has been described for at least nine species of cultured plant cells (Villa and Molinari, 2008). ABA2 is a member of the short-chain alcohol dehydrogenase protein family crucial for ABA biosynthesis (González-Guzmán *et al.*, 2002; Cheng *et al.*, 2002). It catalyzes the conversion of xanthoxin into ABA-aldehyde that is used by the AAO3 protein to produce ABA. Accordingly, the levels of ABA in *aba2* mutants are severely reduced (Léon-Kloosterziel *et al.*, 1996; González-Guzmán *et al.*, 2002), indicating that this is a key step in the basal production of ABA in plants. The conversion of xanthoxin into ABA-aldehyde by ABA2 involves not one but several catalytic steps. ABA2 is able to oxidize the 4’-hydroxyl group of xanthoxin into a 4’-keto group but it also isomerizes the 1’,2’,-epoxide into a 1’-hydroxy-2’ene group (González-Guzmán *et al.*, 2002). Due to structural similarities between xanthoxin and P-Acid B we hypothesized that ABA2 could oxidize the 4’hydroxyl group and dehydrogenate C-2’ and C-3’ of P-Acid B generating ABA. Actually, the enzyme ABA2 can use certain alcohol substrates different from xanthoxin being able to oxidize 3,5,5’-trimethylcyclohexanol alcohol although with higher K_m_ values (González-Guzmán *et al.*, 2002). Taken together we propose P-Acid B as a fungal compound with cross-kingdom activity that might be able to use the plant enzyme ABA2 to induce the synthesis of ABA and inhibit seed germination. A body of evidence indicates that the fungus *P. semeniperda* can benefit from reducing seed germination and seedling establishment through P-Acid B production. *P. semeniperda* reproductive success is often associated with slow germinating or dormant seeds (Finch-Boekweg *et al.*, 2015) while fast germinating seeds usually tolerate *P. semeniperda* infection and develop into normal plants (Finch-Boekweg *et al.*, 2015). Additionally, the distribution of *P. semeniperda* populations is correlated with environmental conditions that reduce seed germination rates and therefore, *P. semeniperda* is found more frequently in dry scenarios rather than humid ones. Dry conditions reduce seed germination and induce postgermination growth arrest (Beckstead *et al.*, 2007, 2016; Meyer *et al.*, 2015).

In summary, we propose that the fungus *P. semeniperda* produces pyrenophoric acid B to inhibit seed germination through the activation of plant ABA biosynthesis, in order to increase its fitness by increasing the probability of pathogen-caused seed mortality, thereby increasing resources available for pathogen reproduction.

## Supplementary data

Fig.S1. Pyrenophoric acid B (P-Acid B) requires ABA2 and onwards to inhibit seedling establishment.

## Acknowledgements

This research was funded in part by Grant JFSP-11-S-2-6 to S.M. from the Joint Fire Sciences Program of the U.S. Departments of Agriculture and Interior and in part, to M.M., by Programme STAR 2017 financially supported by UniNA and Compagnia di San Paolo. J. L.-J. is funded by a Marie-Sklodowska Curie Reintegration Grant H2020-MSCA-707477. Work in P. L. R. lab is supported by RTC-2017-6019-2 and BIO2017-82503-R grants from Ministerio de Ciencia, Innovación y Universidades. M.A.F. is a recipient of a FPU fellowship from MECD. A. Evidente is associated to the Istituto di Chimica Biomolecolare del CNR, Pozzuoli, Italy. We would like to thank Ebe Merilo (University of Tartu) for sharing *nced3,5* and *aba3-1* mutant seeds.

---

Figure S1. **Pyrenophoric acid B (P-Acid B) requires ABA2 and onwards to inhibit seedling establishment.**

(A) Pictures of Arabidopsis seeds treated with 1 μM ABA, 100 μM P-Acid B or DMSO as control, 3 days after sown. (B) Quantification of seedling establishment 3 days after sown seeds on 100 μM P-Acid B. Values represent mean ± SD of 90 seed used in three different experiments.

