## Supplementary figures and images for "The fungal sesquiterpenoid pyrenophoric acid B uses the plant ABA biosynthetic pathway to inhibit seed germination" (from another version of this preprint: version 1, 2019-04-29)

### Supplemental Figure 1

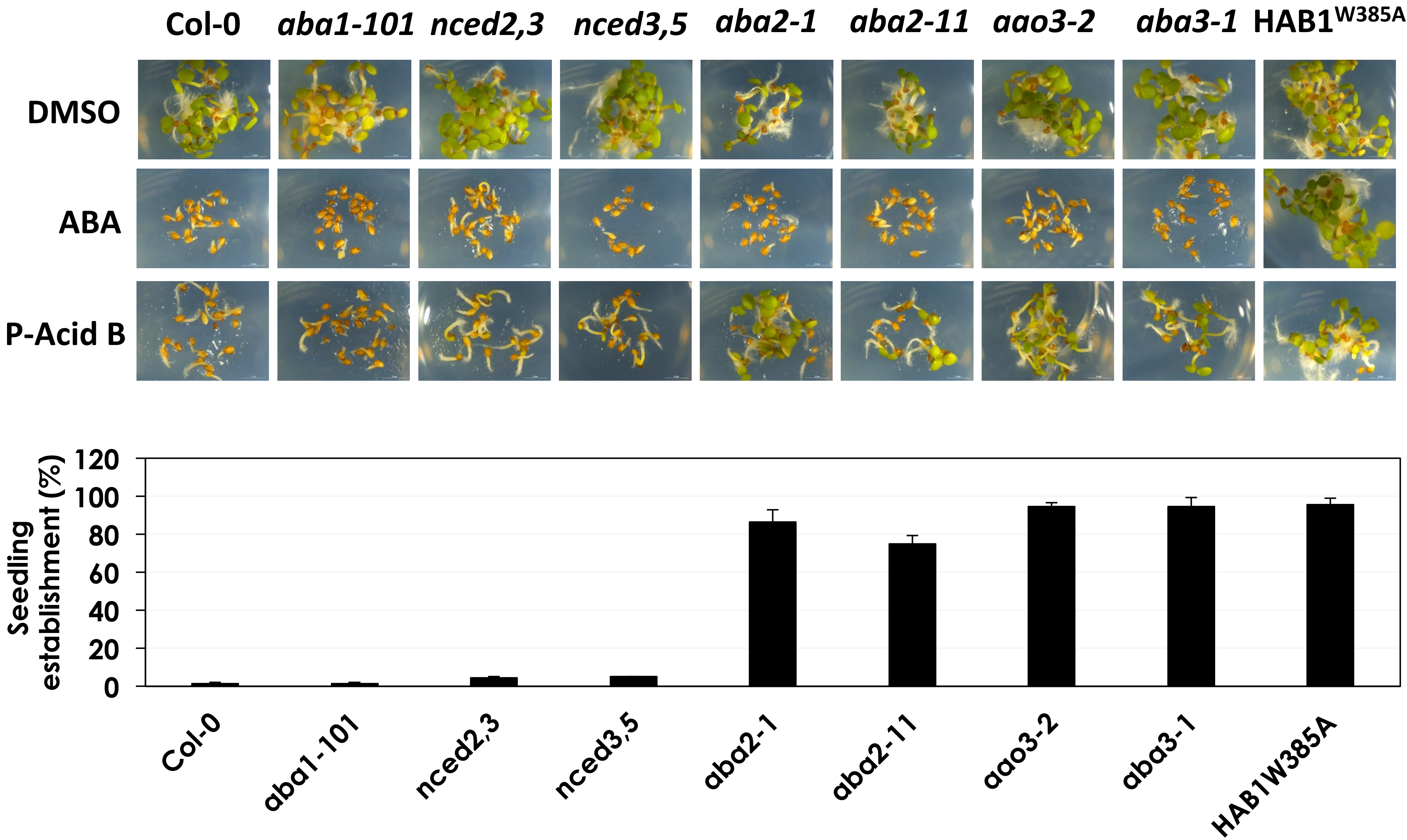
